# PKR-mediated Stress Response Enhances Dengue and Zika Virus Replication

**DOI:** 10.1101/2021.04.08.439069

**Authors:** Taissa Ricciardi-Jorge, Edroaldo Lummertz da Rocha, Edgar Gonzalez-Kozlova, Gabriela Flavia Rodrigues-Luiz, Brian J Ferguson, Trevor Sweeney, Nerea Irigoyen, Daniel Santos Mansur

## Abstract

The mechanisms by which flaviviruses use non-canonical translation to support their replication in host cells are largely unknown. Here we investigated how the integrated stress response (ISR), which promotes translational arrest by eIF2ɑ phosphorylation (p‒eIF2ɑ), regulates flavivirus replication. During Dengue virus (DENV) and Zika virus (ZIKV) infection, eIF2ɑ phosphorylation peaked at 24 hours post infection and was dependent on PKR but not type I interferon. The ISR is activated downstream of p-eIF2α during infection with either virus, but translation arrest only occurred following DENV4 infection. Despite this difference, both DENV4 and ZIKV replication was impaired in cells lacking PKR, independent of IFN-I/NF-kB signalling or cell viability. By using a ZIKV 5′ UTR reporter system as a model, we found that this region of the genome is sufficient to promote an enhancement of viral mRNA translation in the presence of an active ISR. Together we provide evidence that flaviviruses escape ISR translational arrest and co-opt this response to increase viral replication.

## INTRODUCTION

The *Flavivirus* genus is part of the *Flaviviridae* family and contains the causative agents of several human diseases of high global impact, such as Dengue virus (DENV), Zika virus (ZIKV), Japanese encephalitis virus (JEV), yellow fever virus (YFV), West Nile virus (WNV) and others. Flaviviruses are enveloped ssRNA(+) viruses of about 50 nm in diameter with 10-11 kb genome containing a single ORF flanked by two non-coding segments (UTRs) 5′ (capped) and 3′ (non-polyadenylated) (1). Flavivirus replication is cytoplasmic in strong association with ER, forming viral replication complexes by convolution of ER membrane (2).

Despite the presence of a 5′ -cap on its genome, DENV can also use a cap-independent mechanism for protein synthesis under the suppression of the cap-binding protein, eIF4E (3). Using DENV and ZIKV, this mechanism was later shown to be mediated by an internal ribosomal entry site (IRES) localised in viral 5’-UTR (4). A recent report has confirmed the existence of the IRES element (5), indicating, along with previous suggestions (3, 6, 7) that a cap-dependent translation would be used for viral polyprotein synthesis until the suppression of canonical translation initiation when a switch to cap-independent translation would take place. This alternative cap-independent mechanism, including cell conditions to promote the switch in translation mode, remains to be fully characterised.

A well-known cellular control over the translation machinery is coordinated by four kinases of the integrated stress response (ISR), that act as sensors of different stressors. They are: (i) PKR, activated by double-stranded RNA (dsRNA); (ii) HRI, activated by oxidative stress and/or heme deficiency; (iii) PERK, activated by endoplasmic reticulum stress; and (iv) GCN2, activated by deprivation of amino acids (8–10). Therefore, ISR is capable of sensing viral infections directly by detecting replication intermediates (dsRNA) or indirectly *via* structural or metabolic stresses. A shared target of these four kinases is the subunit alpha of the translation factor eIF2 (eIF2α), which is part of the ternary complex (TC, eIF2-GTP-tRNA_i_ ^Met^), necessary for translation initiation. Phosphorylation of eIF2α (p-eIF2α) prevents the exchange of the associated inactive GDP for the active GTP, blocking its recycling for participation in the formation of new 43S complexes, thus inhibiting general protein synthesis (11, 12).

In the present work, we show that PKR is the main kinase leading to eIF2α phosphorylation and cellular protein translation arrest during DENV4 and ZIKV infection. EIF2α phosphorylation, however, is linked to the establishment of an optimal cellular environment for virus replication. Using ZIKV as a model, we demonstrate that the viral 5′-UTR is required to promote enhanced viral translation when eIF2α is phosphorylated. These results suggest that ISR may provide the necessary conditions for viral alternative translation boosting viral replication.

## RESULTS

### DENV4 and ZIKV induce ISR activation during infection

DENV4 and ZIKV infections were carried out in human lung A549 cells. These cells support viral replication and respond to type one interferons (IFN-I) (4, 13–17). Interferon stimulated genes (ISGs) are central for flavivirus restriction. We evaluated eIF2α phosphorylation during DENV4 and ZIKV infection as an ISR activation marker. By immunoblot analysis, we showed that both viruses induce p‒eIF2α by 24 hours post infection (h p.i.) (Figure 1A). Furthermore, this time point coincides with the detection of p-PKR (Thr451), the activated form of the protein, suggesting the participation of this kinase in ISR activation during infection. Quantification by flow cytometry enabled us to determine that eIF2α phosphorylation happened after 12 h p.i. and peaking at 24 h p.i. (Figure 1B). This time frame is surprisingly late, considering that from at least 12 h p.i. infectious virus particles can already be detected in cell supernatants under these experimental conditions (Supplementary Figure 1).

**Figure 1.**
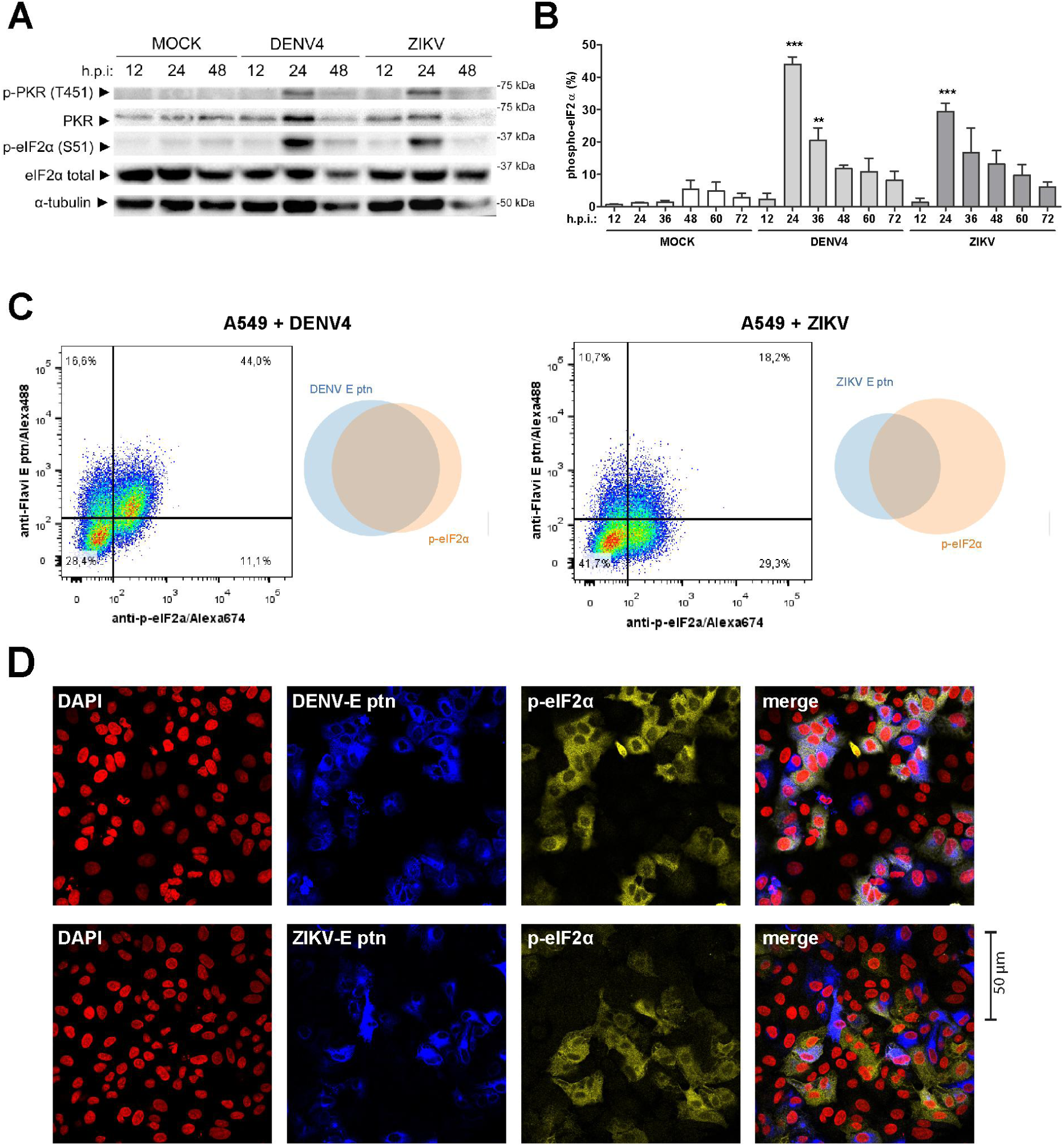
DENV4 and ZIKV promote phosphorylation of eIF2α in A549 cells. A549 cells were infected with DENV4 (MOI 2) or ZIKV (MOI 3) and harvested at the indicated times for analysis. (A) Immunoblot analysis of cell extracts resolved in denaturing SDS-PAGE. (B) Flow cytometry analysis for quantification of cells expressing p‒eIF2α within the total population (C) Dot plot and Venn diagram of representative results from co-staining with anti-flavivirus E protein and anti-p‒eIF2α at 24 h p.i. analysed by flow cytometry and (D) immunofluorescence of infections under the same conditions, representative images of three independent experiments. In the column chart, the bars represent the means ± SEM from three independent experiments. Statistical analysis was performed by paired *t* test comparing infected samples to the respective uninfected control. *: p ≤0.05; **: p ≤0.01; ***: p ≤0.001; ns / no markup: no statistical difference; h p.i: hours post-infection.

To further investigate the context of eIF2α phosphorylation, cells were co-stained with a monoclonal antibody directed against the flavivirus envelope protein (Figure 1C-D). At 24 h p.i. DENV4 infected cells exhibited a significant double-positive population, with about 90% of p‒eIF2α+ cells being infected. However, during ZIKV infection, there is a lower proportion of double-positive cells, with about 40% of envelope positive cells in the p‒eIF2α+ population. These cells may be infected with ZIKV, but they do not yet express enough envelope protein to allow detectable staining. The same profile of co-staining quantified by cytometry could also be visualised by immunofluorescence (Figure 1D).

In summary, although both infections led to different profiles of anti‒p‒eIF2α and anti‒flavivirus co-staining, both viruses promoted stress responses within the same time frame.

### Phosphorylation of eIF2α during DENV4/ZIKV infection is PKR-dependent and IFN-I independent

To investigate which ISR kinase promotes eIF2α phosphorylation in our model, gene deletion by CRISPR-Cas9 was used to produce A549 PKR and IFNAR knockout cell lines (Supplementary Figure 2A). In PKR^-/-^ cells infected with DENV4 or ZIKV, p‒eIF2α was abrogated indicating that ISR activation promoted by both viruses is mostly dependent on this kinase (Figure 2A,B). Since PKR is an IFN-stimulated gene (ISG), and considering the late ISR activation, the participation of IFN-I signalling in eIF2α phosphorylation was assessed. However, there was no change in the level of p‒eIF2α in infected IFNAR^-/-^ A549s compared to WT cells (Figure 2A,B). Likewise, experiments using a PKR/IFNAR double-knockout (Supplementary Figure 2B) lineage confirmed the requirement for constitutive PKR expression to drive eIF2α phosphorylation.

**Figure 2.**
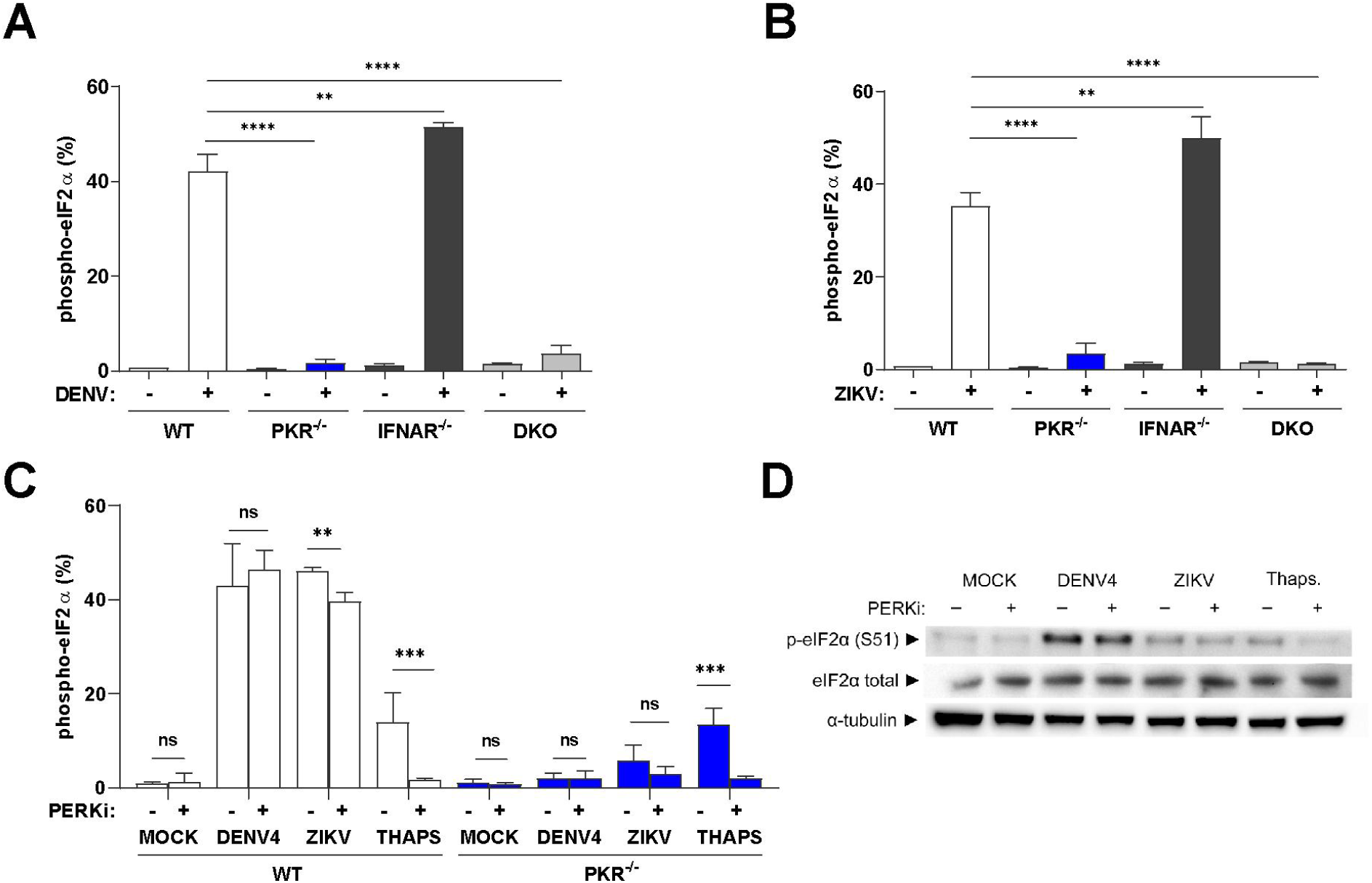
Phosphorylation of eIF2α during DENV4 or ZIKV infections is PKR-dependent and IFN-independent. (A) Flow cytometry analysis for quantification of cells expressing p‒eIF2α within the total population of the indicated lineages of A549 cells infected with DENV4 (MOI 2) at 24 h p.i.; and (B) same experiment using ZIKV (MOI 3) (C) Flow cytometry analysis for quantification of cells expressing p‒eIF2α within the total population A549 WT and PKR^-/-^cells infected with DENV4 (MOI 2) or ZIKV (MOI 3) or treated with UPR inducer thapsigargin 2µM for 45 minutes in the presence or absence of the PERK inhibitor GSK2656157 (5 µM) for 24h; and (D) immunoblot analysis of cell extracts from the same experiment resolved in denaturing SDS-PAGE. Representative image of two independent experiments. DKO: double IFNAR^-/-^/PKR^-/-^ knockout. In column charts, bars represent the mean ± SEM from three independent experiments. Statistical analysis was performed by one-way ANOVA, followed by Tukey’s test for multiple comparisons. *: p ≤0.05; **: p ≤0.01; ***: p ≤0.001; ns / no markup: no statistical difference.

The persistence of a small population p‒eIF2α+ in the PKR^-/-^ cells infected with ZIKV, indicates the activation of other ISR kinase by this virus. Therefore, we decided to investigate PERK activity using the selective inhibitor, GSK2656157 (18). GSK2656157 (PERKi) treatment did not alter the induction of p‒eIF2α in DENV4 infected cells but promoted a small decrease in p‒eIF2α population in ZIKV infected cells (Figure 2C). This demonstrated a minor participation of PERK in the ISR activation in the ZIKV model.

In summary, the phosphorylation of eIF2α during DENV4 or ZIKV infection in this model was mainly PKR dependent and IFN-I independent.

### PKR is not blocked by DENV4 or ZIKV but shows delayed activation

The late PKR activation during infection with both viruses is not explained by the need of type I IFN signalling to boost PKR levels. Therefore, we tested whether PKR activity is blocked by infection at earlier time points. To test this, the dsRNA analogue, poly (I:C), was used to stimulate PKR. Poly (I:C) transfection was performed at 3 h p.i. and cells were analysed at 9 h p.i., that is, before virus-promoted eIF2α phosphorylation (Figure 3A). The comparable levels of p‒eIF2α between mock and infected cells stimulated with poly (I:C) suggest that PKR is not blocked by DENV4 or ZIKV in the early stages of infection (Figure 3A). The absence of p‒eIF2α in PKR^-/-^ cells demonstrates that eIF2α phosphorylation induced by poly (I:C) is exclusively dependent on this kinase.

**Figure 3.**
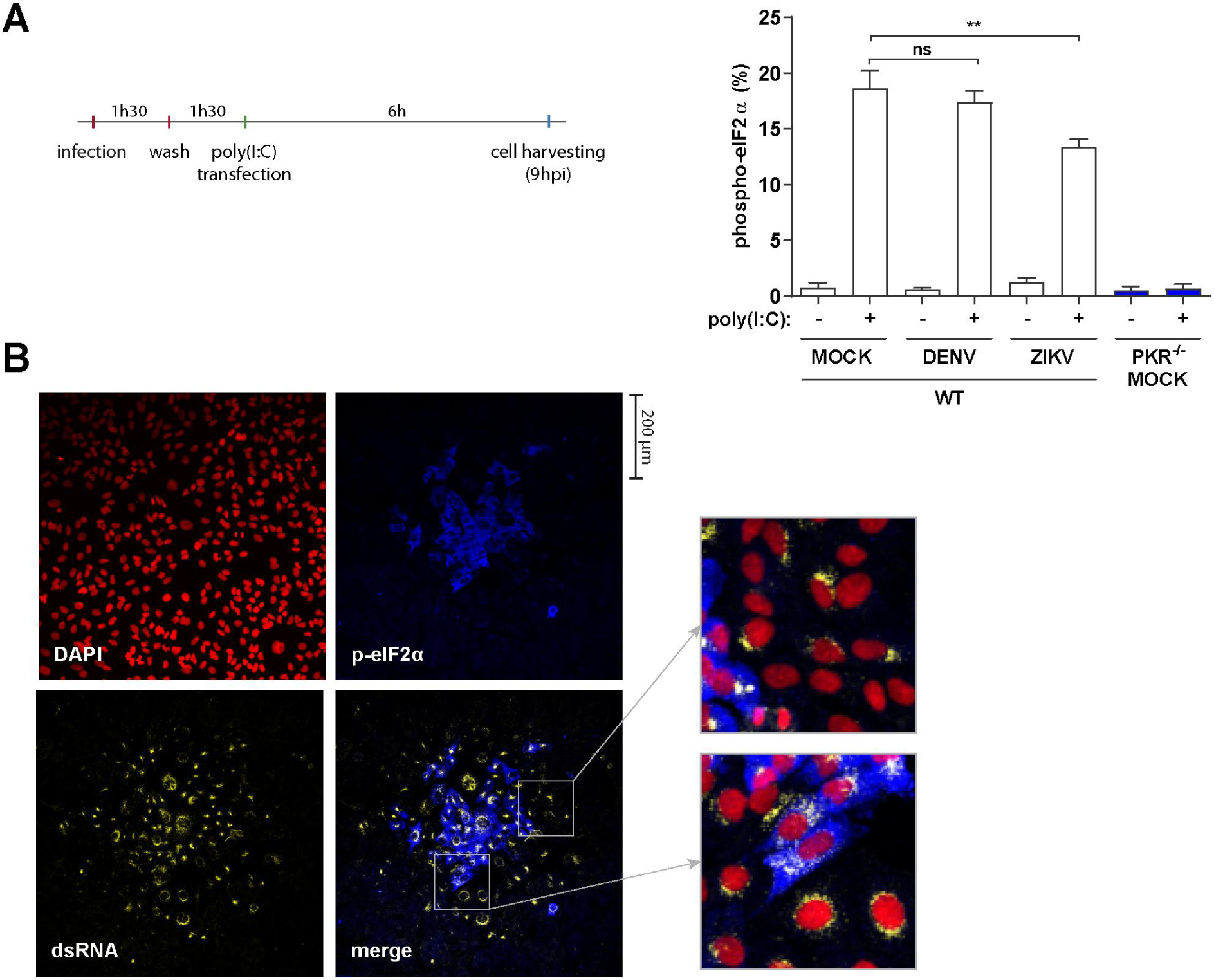
PKR is not blocked by DENV4 or ZIKV but shows delayed activation. (A) Cells were infected with DENV4 (MOI 2) and ZIKV (MOI 3) and incubated for 1h 30min before the virus inoculum was removed. Then, cells were stimulated with poly (I:C) at 3 h p.i. with 10 μg/ml for 6 hours. Analysis for quantification of the cell population expressing p‒eIF2α after stimulation with poly (I:C) was performed by flow cytometry. Data from three independent experiments. (B) Immunofluorescence of A549 WT cells infected with DENV4 for 48 h p.i. in semisolid medium. Cells fixed, permeabilized and co-stained with DAPI, anti-dsRNA and anti-p‒eIF2α. Representative image of three independent experiments. In the column chart, bars represent the means ± SEM. Statistical analysis was performed by one-way ANOVA, followed by Tukey’s test for multiple comparisons. **: p ≤0.01; ns: no statistical difference.

If PKR is not blocked in the early stages of infection by DENV4 or ZIKV, the late ISR activation could be due to the amount of dsRNA in infected cells not reaching the threshold required for PKR activation. To test this hypothesis, infected cells were incubated in a semisolid medium so viral spread happens from cell to cell by juxtaposition allowing temporal inferences of events based on cell location from the centre of the viral foci. Immunofluorescence of DENV4 infected cells shows that cells at the edge of the viral foci do not phosphorylate eIF2α despite the presence of large amounts of dsRNA (Figure 3B). This delayed activation of PKR may be explained by the ability of flaviviruses to hide their dsRNA inside viral replication complexes, as suggested for other RNA sensors (19–21). Another possibility is PKR being activated not by viral dsRNA, but by mitochondrial RNA as a consequence of viral suppression of mitophagy (22). In either case, these results indicate that at this stage of the infection, ISR activation by PKR is delayed, occurring in infected cells after the establishment of viral replication complexes. Further studies will be necessary to determine the nature of this delay.

### DENV4, but not ZIKV, induces translation arrest in infected cells

To investigate the consequences of eIF2α phosphorylation on cellular and viral protein synthesis, viral infections were performed in semi-solid medium and a puromycin labelling protocol was used to assess general translation activity (23). By immunofluorescence analysis, it was possible to observe that p‒eIF2α in WT lineage promotes cellular translation arrest during DENV4 infection, evidenced by the absence of puromycin incorporation in the centre of viral plaques (Figure 4A). At the edges of the plaque, however, anti-puromycin and anti-flavivirus simultaneous staining reinforces data presented previously (Fig 3B) that translation arrest happens relatively late during DENV4 infection. In PKR^-/-^ cells, however, co-staining is present throughout the viral plaque, suggesting cell translation downregulation during DENV4 infection is caused by ISR activation via PKR.

**Figure 4.**
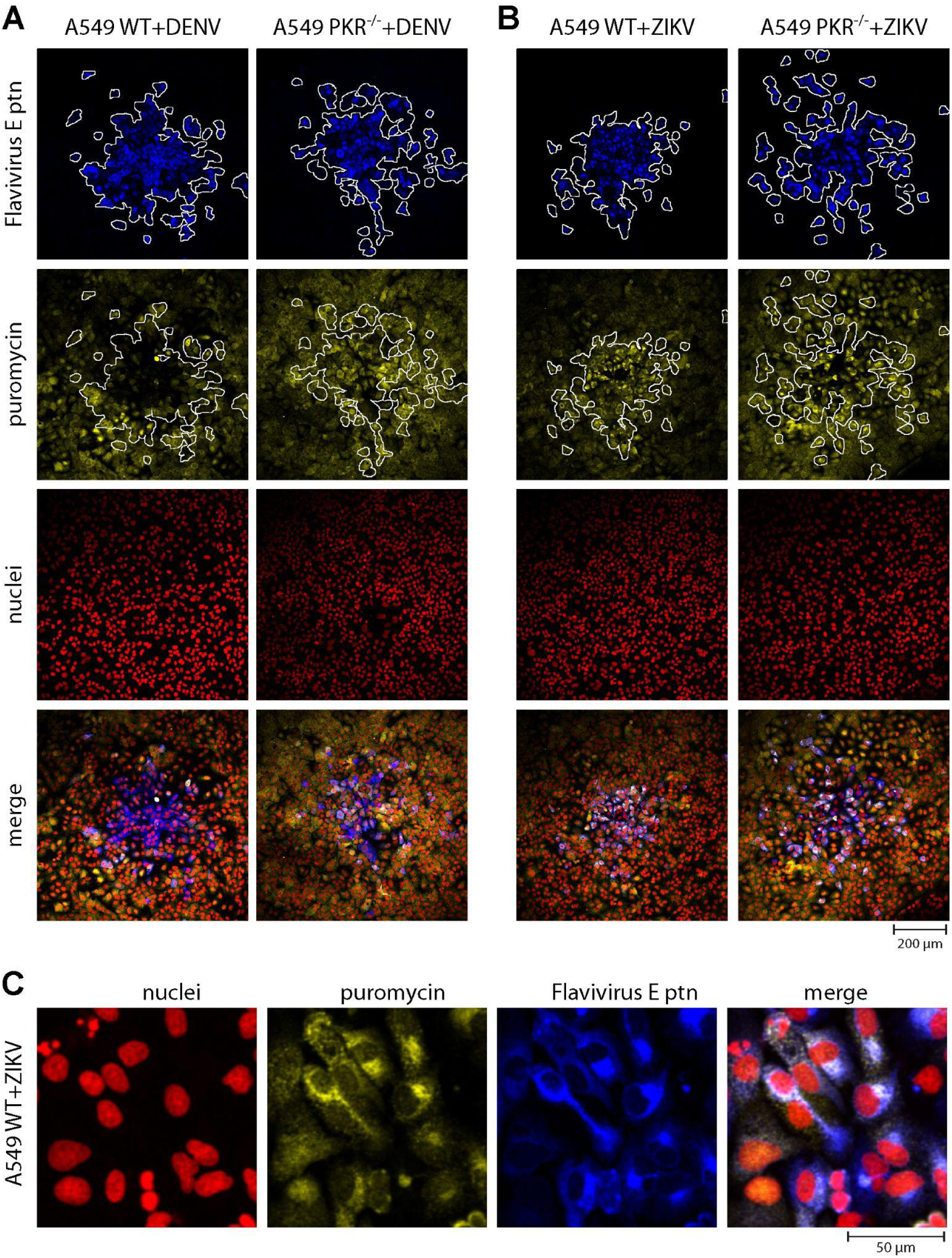
DENV4, but not ZIKV, induces PKR-dependent translation arrest in infected cells. Immunofluorescence of A549 WT or PKR^-/-^ cells infected with DENV4 (A) or ZIKV (B) for 48 h p.i. in semisolid medium. Cells labelled with puromycin before fixation and permeabilization. Co-staining with DAPI, anti-puromycin, anti-flavivirus envelope (E) and secondary antibodies. (C) Zoom on WT cells infected with ZIKV for a better appreciation of the immunostaining pattern. Representative images of three different experiments.

Surprisingly for ZIKV, there is a different profile, with more intense incorporation of puromycin by infected WT cells than bystander cells (Figure 4B). Bystander cells are here defined as cells that were in the same environment of infected cells (i.e., same well or organism) but viral protein was not detected. Anti-puromycin signal is stronger in the perinuclear region which coincides with viral replication sites (Figure 4C). The phenotype of increased translation activity in infected cells is lost in PKR^-/-^cell lines, suggesting that PKR activation allows increased viral translation.

### PKR is required for expression of eIF2α-downstream genes and proteins during DENV4 and ZIKV infections

Phosphorylation of eIF2α as indicated above results in a general shutdown of protein synthesis, however, translation of the activation transcription factor 4 (ATF4) is increased in this situation (24, 25) leading to the induction of its target genes *DDIT3 (CHOP)* and *GADD34* involved in cellular stress resolution pathways (26, 27)GADD34 is responsible for eIF2α de-phosphorylation hence resuming global translation. Thus, to evaluate the impact of PKR deficiency in events downstream of eIF2α phosphorylation, the expression of *DDIT3* (CHOP) and *GADD34* was analysed by qPCR. As observed in Figure 5A, both *DDIT3* and *GADD34* were less expressed in PKR^-/-^ cells than in WT cells during DENV4 and ZIKV infection. This downregulation is subtle, possibly due to control of these genes by other pathways such as IRF3/7 (28, 29) and STAT3-NFkB or NF-YA (30, 31). Consistent with its regulation by eIF2:p-eIF2 levels (32, 33), when GADD34 protein is analysed by immunoblotting (Figure 5B) or flow cytometry (Figure 5C), this protein shows a pattern of expression similar to that of p‒eIF2α in WT and PKR^-/-^ cells (Fig. 2A-B), demonstrating the transcriptional and post-transcriptional cellular control over eIF2α downstream signalling.

**Figure 5.**
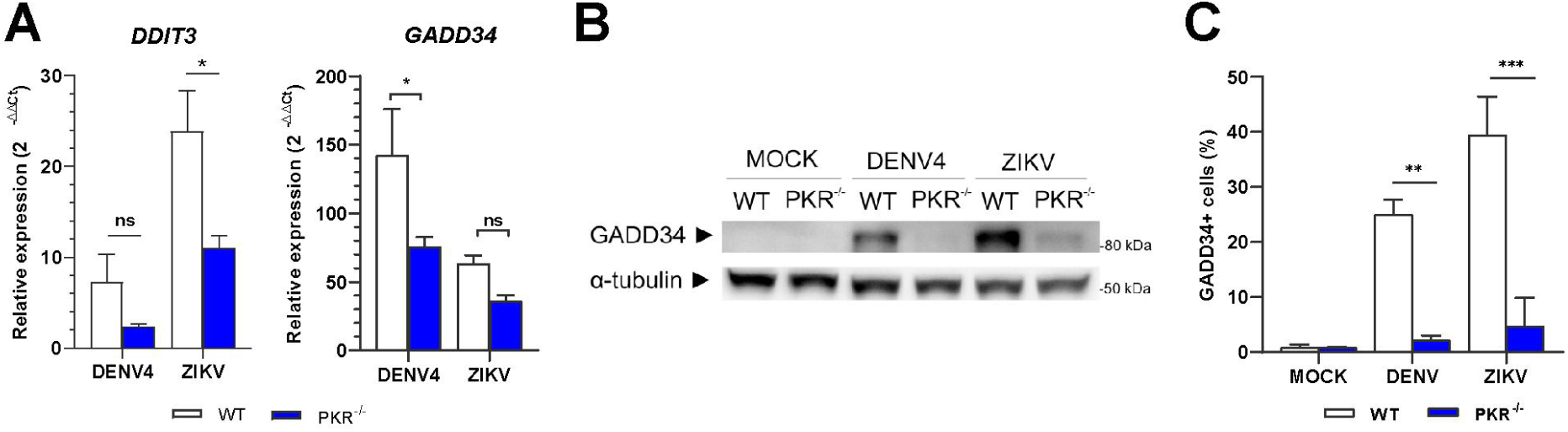
PKR is required for expression of eIF2α-downstream genes and proteins during DENV4/ZIKV infections. A549 WT or PKR-/-cells infected with DENV4 (MOI 2) or ZIKV (MOI 3) and harvested at 24 h p.i. for analysis. (A) Quantification by RT-qPCR of DDIT3 and GADD34 gene expression. Relative expression calculated by 2-ΔΔCt methods using mock cells as reference. (B) Immunoblot analysis of GADD34 protein expression. Representative image of two independent experiments. (C) Flow cytometry analysis for quantification of cells expressing GADD34 within the total population. In the column charts, bars represent the means ± SEM from three independent experiments. Statistical analysis was performed by paired *t* test comparing the two cell lineages under the same conditions. *: p ≤0.05; **: p ≤0.01; ***: p ≤0.001; ns / no markup: no statistical difference.

Together, these results show that during infection by DENV4 and ZIKV, the activation of the stress response is not restricted to phosphorylation of PKR and eIF2α; and reinforces that the activation of this pathway is abolished in the PKR^-/-^ cells.

### Disruption of the PKR‒eIF2α pathway impairs DENV4 and ZIKV replication

Considering the efficacy of p-eIF2α in blocking cellular translation and the dependency of PKR for this phosphorylation, we evaluated the impact of PKR deletion on virus replication. Surprisingly, replication of both viruses was impaired in PKR^-/-^cells, shown by lower virus titres (Figure 6A) and smaller plaque sizes and number of cells/plaque (Figure 6B-C). Viral genome replication, assessed by qPCR, does not show statistically significant difference, but a trend of lower replication in PKR^-/-^ cells can be seen and may reflect impairment of other viral replication steps (Figure 6D).

**Figure 6.**
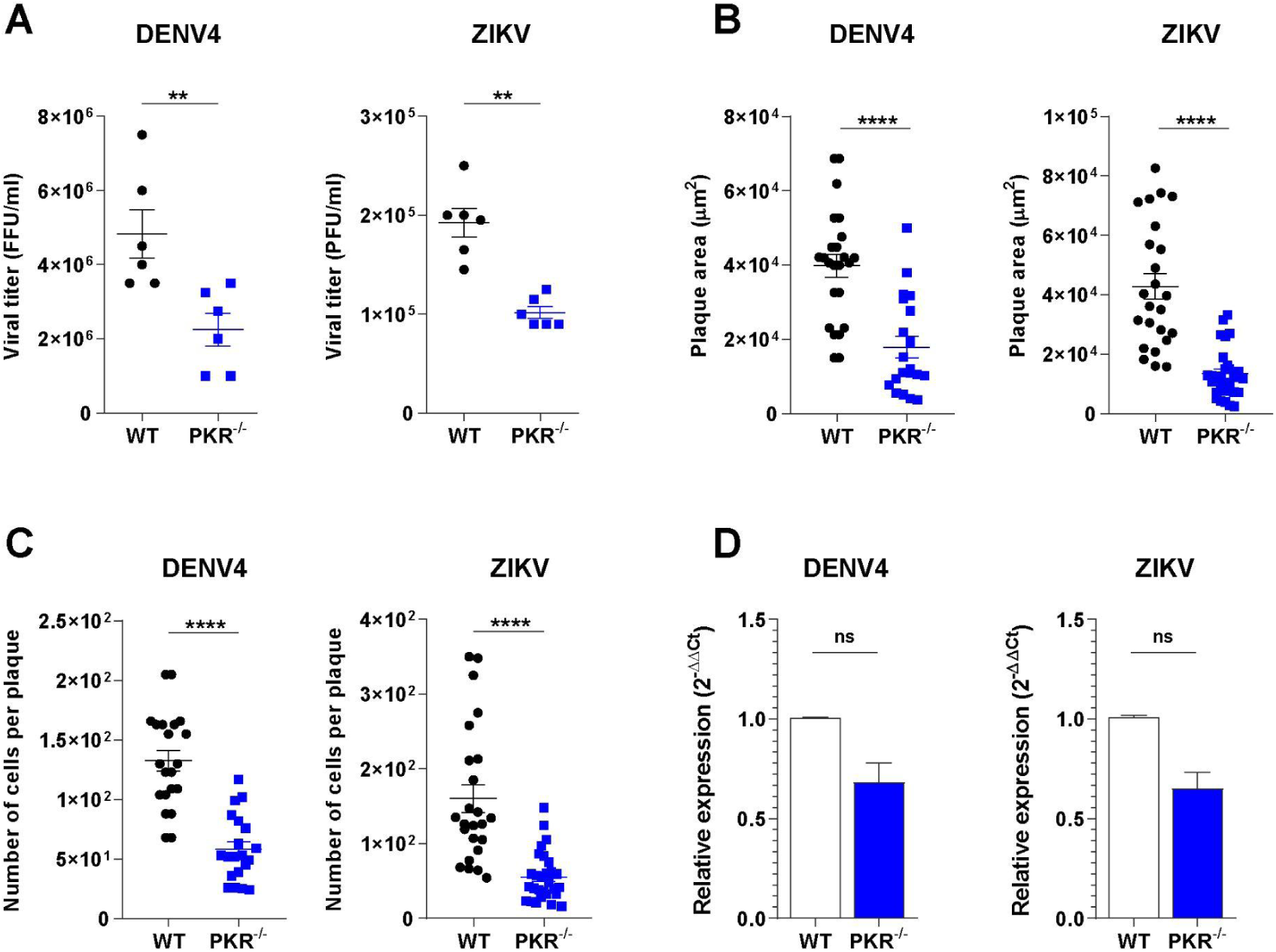
Disruption of the PKR‒eIF2α pathway impairs DENV4 and ZIKV replication. (A) Viral titration from A549 WT or PKR^-/-^ cells infected with DENV4 (MOI 2) or ZIKV (MOI 3) and harvested at 24 h p.i. for analysis. Plaque size comparison of virus in A549 WT or PKR^-/-^ cells infected with DENV4 and incubated for 8 days or infected with ZIKV and incubated for 5 days: measurements of (B) plaque areas and (C) number of cells per plaque from immunofluorescence. (D) Quantification by RT-qPCR of DENV4 or ZIKV viral RNA from A549 WT or PKR^-/-^ cells infected with DENV4 (MOI 2) or ZIKV (MOI 3) and harvested at 24 h p.i. for analysis. Relative expression calculated by 2^-ΔΔCt^ method using mock cells as a reference for cell genes and WT cells as reference for viral genome. PFU: plate forming units; FFU: focus forming units. In all charts, bars represent the means ± SEM from three independent experiments. Statistical analysis was performed by paired *t* test comparing the two cell lineages under the same conditions. *: p ≤0.05; **: p ≤0.01; ***: p ≤0.001; ns / no markup: no statistical difference.

The lower virus titres and diminished plaque sizes in PKR^-/-^ cells are particularly counterintuitive since this cell line is defective in activating the whole antiviral pathway coordinated by PKR. This finding indicated the possibility that the PKR deletion has a negative impact on other antiviral pathways and/or these viruses could have co-opted the ISR to promote viral replication.

### PKR deletion does not affect the innate immune response or cell viability

The phenotype of reduced viral replication in PKR^-/-^ lineage could be explained by changes in the cellular innate immune response, as several studies report the role of PKR in sustaining NF-κB and IFN signalling (34–39) and promoting IFNγ and TNFα expression (40, 41). To investigate the possibility of alterations in innate immune pathways as a result of PKR deletion, the gene and protein expression of elements of this cellular response were evaluated.

Genes activated by IRF3/7, IFN-I or NF-κB did not show any significant difference in transcription between WT and PKR^-/-^ cells during virus infections (Figure 7A). Immunoblotting corroborated this transcriptional analysis, as no differences were observed in STAT1 activation, or IFIT1 and ISG15 expression levels between the two cell lines, thus demonstrating that virus-induced IFN-I signalling was not affected by PKR deletion (Figure 7B).

**Figure 7.**
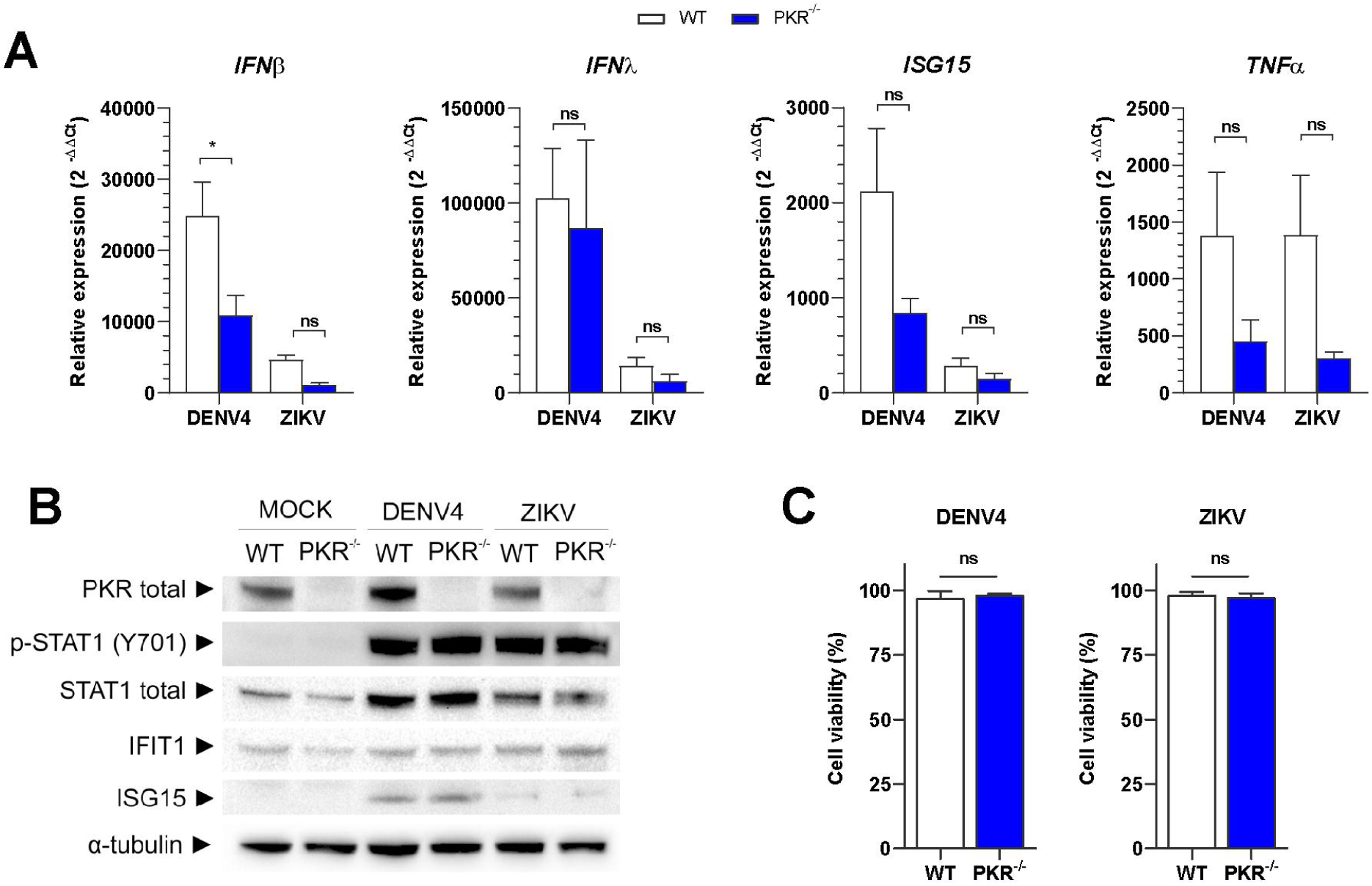
PKR deletion does not affect the innate immune response or cell viability. A549 WT or PKR-/-cells infected with DENV4 (MOI 2) or ZIKV (MOI 3) and harvested at 24 h p.i. for analysis. (A) Quantification by RT-qPCR of *IFNβ*, *IFNλ*, *ISG15* and *TNFα* gene expression. Relative expression calculated by 2^-ΔΔCt^ methods using mock cells as a reference. (B) Immunoblot analysis of cell extracts resolved in denaturing SDS-PAGE. Representative image of two independent experiments. (C) Flow cytometry analysis for quantification of living cells by staining with Zombie-NIR viability dye. Mock WT cells set as 100% reference. In the column charts, bars represent the means ± SEM from three independent experiments. Statistical analysis was performed by paired *t* test comparing the two cell lineages under the same conditions. *: p ≤0.05; **: p ≤0.01; ***: p ≤0.001; ns / no markup: no statistical difference.

As PKR could also play a role in cell survival during viral infection (42), cell viability was analysed at 24 h p.i. by flow-cytometry. Also in this assay, no difference between WT and PKR^-/-^ cells was observed during virus infections (Figure 7C).

These results allow us to conclude that PKR deletion does not significantly affect the innate immune response or cell survival during DENV4 or ZIKV infection. Therefore, alterations in these pathways do not explain the lower viral replication seen in the PKR^-/-^ cells.

### ZIKV co-opts PKR‒eIF2α pathway to increase viral translation through its 5′ -UTR

ISR control over cell translation can promote the expression of certain mRNAs as a consequence of their 5′ -UTR sequence and/or structure (33, 43). Therefore, we next sought to investigate whether viral 5′ -UTR could subvert the ISR to promote replication. Hence, a translation activity assay was performed based on the ZIKV genome using a luciferase reporter readout. For this assay, a reporter plasmid was designed with the first 194 nucleotides of ZIKV, this includes the 5′ -UTR followed by the first 87 nt from the viral polyprotein fused in frame with the firefly luciferase gene (Figure 8A). A control plasmid for expression of *Renilla* luciferase with the same backbone that the firefly luciferase reporter was co-transfected for data normalisation. Results are presented in Relative Translation Activity (RTA), given in percentage, using values obtained with WT mock cells as reference (100%).

**Figure 8.**
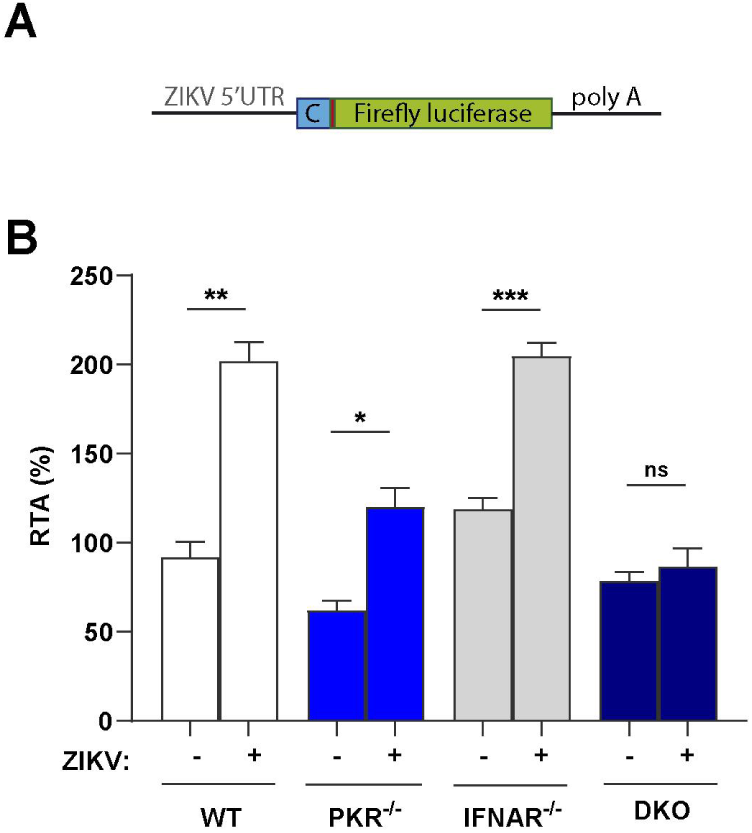
ZIKV 5′ -UTR is sufficient to increase relative reporter translation in PKR‒eIF2α competent cells. (A) Diagram of reporter construct used for the assay, containing the first 87 nt of viral capsid protein. (B) Relative translation of ZIKV reporter in A549 (WT/PKR^-/-^/IFNAR^-/-^/DKO) cells infected with ZIKV (MOI 3), transfected at 12 h p.i. and analysed at 36 h p.i. In the column chart, the bars represent the means ± SEM from three independent experiments. *: p ≤0.05; **: p ≤0.01; ***: p ≤0.001; ns / no markup: no statistical difference.

Cells were infected with ZIKV (or MOCK treated) for 12 hours before plasmid transfection, and then analysed at 36 h p.i. to maximise the time of luciferase expression under the context of ISR activation. The comparison between mock and infected conditions for each cell line reveals that virus infection promotes a 2-fold increase in translation activity by WT and IFNAR^-/-^cells (Figure 8B). In contrast, in PKR^-/-^ and double knockout cells this effect is suppressed and the relative translation activity in these cells is similar or lower to WT uninfected cells. These results suggest that the ZIKV 5′ -UTR is sufficient to promote a relative advantage of mRNA translation in the presence of p‒eIF2α.

## DISCUSSION

Flaviviruses efficiently translate their genome via a cap-dependent translation mechanism. However, following suppression of canonical translation initiation that takes place during infection, it has been proposed that a switch to cap-independent translation occurs (3, 5, 44). Cell conditions to promote the switch in translation mode and the required auxiliary factors (e.g. eIFs) for alternative translation remain to be fully characterised.

In the present work, we investigated how the ISR, known to promote translation arrest through eIF2α phosphorylation, would play a role in this elusive non-canonical translation mechanism employed by flaviviruses. Using A549 cells we verified that PKR promotes p-eIF2α during DENV4 and ZIKV infections. In accordance, previous reports have also detected p-eIF2α in A549 cells during DENV (6, 45), ZIKV (46) and JEV (47) infections; as well as studies using other cell lines (48, 49). Further characterization of p-eIF2α was performed by flow cytometry to allow quantification and co-staining for other markers. Although DENV4 and ZIKV promote eIF2α phosphorylation within the same time frame, these viruses displayed different profiles of anti‒p‒eIF2α and anti‒flavivirus envelope protein co-staining, indicating distinct dynamics between viral replication and activation of stress response for each virus. Therefore, cytometry is here proposed as a complementary method to investigate ISR activation, in addition to immunoblotting and immunofluorescence, as it provides more detailed information.

It is worth noting that a few studies observe the activation of the PERK pathway of the unfolded protein response (UPR) during DENV and ZIKV infections without excluding the participation of other ISR kinases especially regarding eIF2α phosphorylation (50–54). In the present model, PERK plays a minor role in ZIKV-induced p-eIF2α. Here, PKR is clearly accountable for the majority of ISR activation during infection with both DENV4 and ZIKV. The ISR was shown to be activated by PKR to downstream genes via eIF2α phosphorylation in WT cells but abolished in PKR^-/-^ cells. However, other flaviviruses such as JEV and WNV seem to phosphorylate eIF2α at least partially dependent on PERK (50–54). For JEV, this could be due to PKR blocking promoted by viral NS2a (47).

An increased expression of p-eIF2α downstream genes was also found in Huh7 cells infected with DENV (55). In this study, single-cell RNA-seq analysis showed upregulation of *DDIT3* and *GADD34* (aka *PPP1R15A*) mRNAs, as well as of other ATF4-downstream genes (*ASNS, CTH, HERPUD1, SOD2, TRIB3*) (56–60) within the infected population (Supplementary Figure 3A-B). A second study using the same approach in PBMC (61) revealed an upregulation of *GADD34* mRNA in DENV-infected monocytes and T cells (Supplementary Figure 3C-D). The upregulation of these mRNAs suggests ISR activation after DENV infection in these cell models. However, as shown here, the expression of these genes is also regulated by post-transcriptional mechanisms dependent on the presence of p-eIF2α.

In our model, a PKR-dependent translation arrest was confirmed in DENV-infected cells as expected, but curiously not in ZIKV-infected cells. For the latter, increased puromycin incorporation in perinuclear areas indicates a possible viral escape mechanism from ISR translation control. This difference might be related to the distinct dynamics between viral replication and activation of stress response for each virus seen in the co-staining assay. Therefore, it calls for further investigation using methods such as live-cell imaging, to track down the sequence of events during infection.

Roth et al, reported a general translation arrest after infection of Huh7 cells with several *Flavivirus* species and strains despite the absence of p-eIF2α (7). This conflicting result could be due to the different cells used. Although we do not exclude the possibility that other translation suppression mechanisms, such as RIDD (62), RNase L (63) and 4E-BP dephosphorylation (64, 65), can also take place in our model in later times after infection.

Despite the differences in translational activity under activated ISR, both DENV4 and ZIKV presented an approximated 50% less viral replication in PKR^-/-^ cells. This decrease suggests that ISR activation is not essential for viral replication but rather exploited by both viruses to counter a major cellular antiviral pathway for their own benefit. Indeed, after discarding the hypothesis of PKR deletion disturbing common antiviral pathways in this model (e.g., type I IFN expression, NFκB activation and apoptosis), our translation activity assay has shown that the ZIKV 5′ -UTR promotes a relative advantage of mRNA translation in the presence of p‒eIF2α. This finding reveals that ZIKV not only escapes ISR translation arrest, but also uses this response to increase viral replication. We hypothesise that this could be achieved by the presence of multiple upstream open reading frames (uORFs) in the 5′ region of the ZIKV genome (66). The presence of multiple uORFs is what allows the main ORF of ATF4 to be translated in the presence of p-eIF2α (24, 25), therefore, a similar strategy could be used by ZIKV to translate the viral polyprotein during ISR activation.

However, we can not rule out that viral alternative translation that does not require eIF2/TC for initiation is also used by DENV4 and ZIKV. The independence of eIF2 for translation initiation under stress conditions has been previously reported within *Flaviviridae* family, since HCV (*Hepacivirus* genus) and *Pestivirus C* (formerly CSFV, *Pestivirus* genus) IRES present similar functioning through IRES dependent mechanisms (67–69).

Here we demonstrate that DENV4 and ZIKV induce eIF2α phosphorylation in a PKR dependent manner and that its absence is detrimental for optimal virus replication. Using ZIKV as a model we showed that this effect maps to the 5′-UTR of the viral genome. Altogether these results suggest that the activation of the PKR/p-eIF2α axis may be necessary for the engagement of the non-canonical translation employed by flaviviruses.

---

**Supplementary Figure 1.**
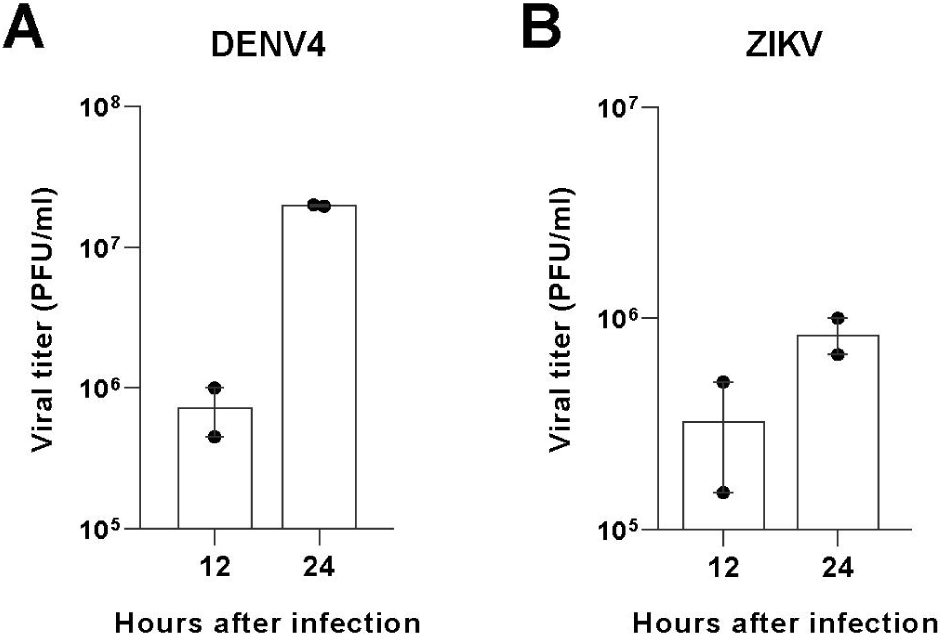
Viral titration in VERO cells of supernatant from A549 WT cells infected with (A) DENV4 (MOI 2) or (B) ZIKV (MOI 3) harvested at 12 and 24 h p.i. for analysis. PFU: plate forming units; FFU: focus forming units. In all charts, points represent the means ± SEM from three independent experiments.

**Supplementary Figure 2.**
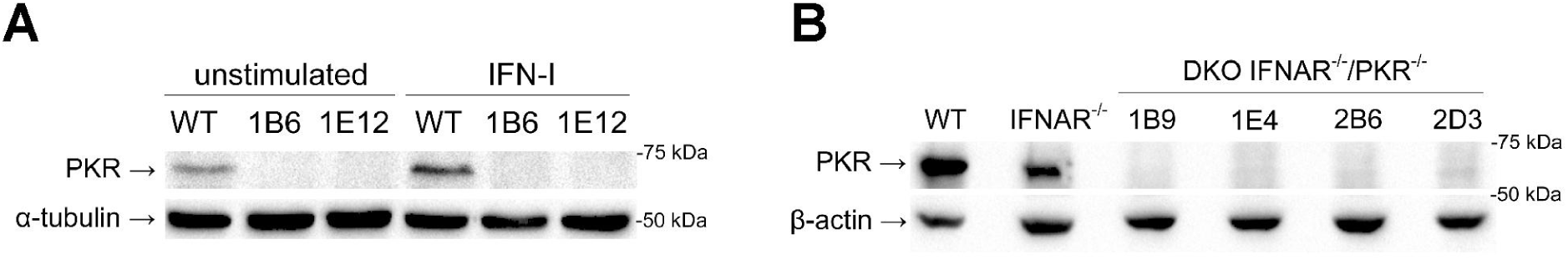
Characterization of knockout cell lines. (A) Clones of A549 PKR^-/-^: Immunoblot of unstimulated and stimulated cells with 100 IU/ml of IFN-α2a for 12 hours demonstrating successful PKR deletion. (B) Clones of A549 IFNAR^-/-^/PKR^-/-^ after stimulation with 100 IU/ml of IFN-α2a for 12 hours demonstrating the deletion of PKR on the parental IFNAR^-/-^ cells. DKO: double-knockout.

**Supplementary figure 3.**
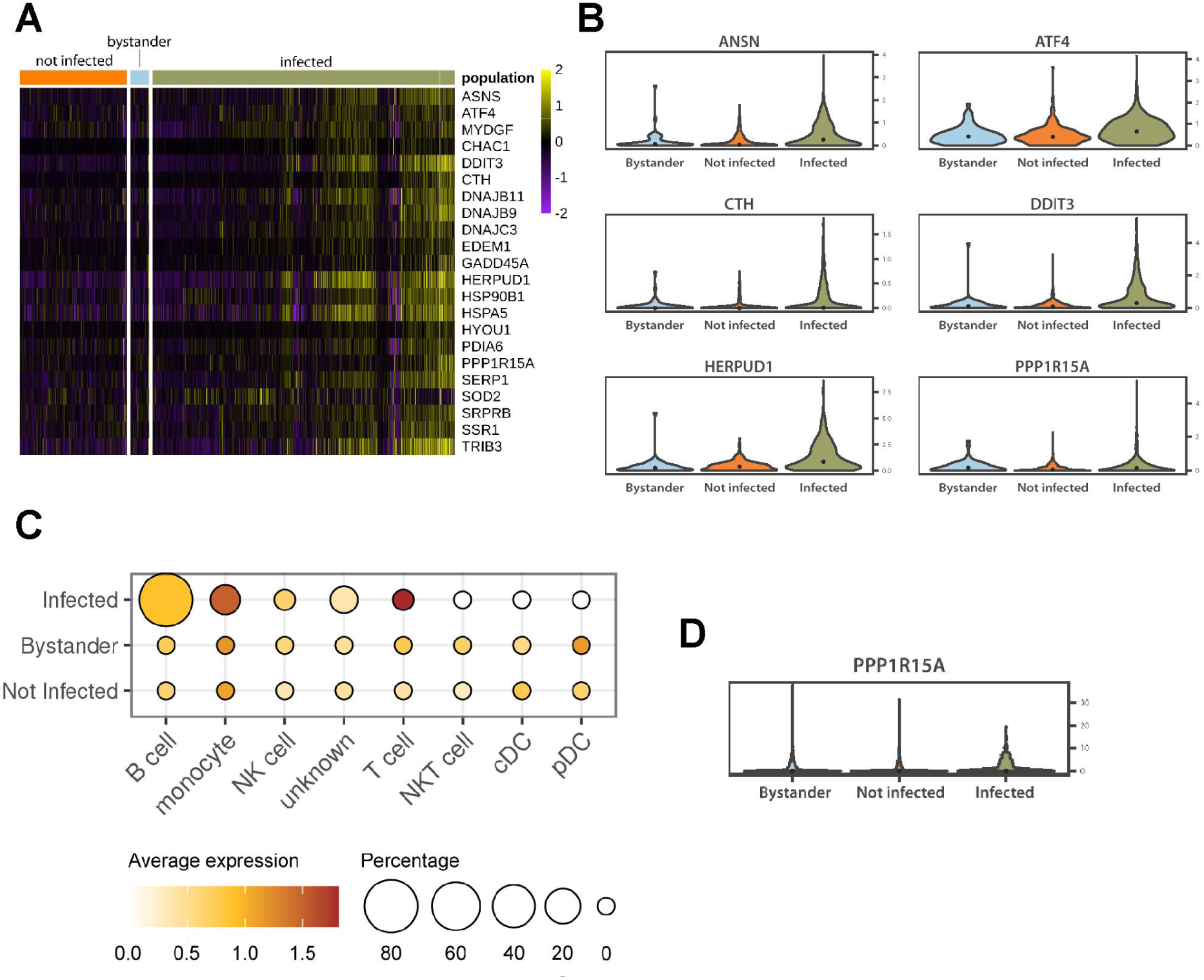
p‒eIF2α-downstream genes are upregulated in DENV-infected Huh7 cells and PBMCs. (A) Dataset from (55), of DENV-infected Huh7 cells, analysed by single-cell RNA-seq. Heatmap representing expression variability of stress-related genes in uninfected, bystander and infected cells; and (B) violin plot representing the expression of p-eIF2α and ATF4-downstream genes in uninfected, bystander and infected cell populations. (C) Dataset from (61), of PBMCs from patients infected with DENV, analysed by single-cell RNA-seq. Dot plot representing the expression of PPP1R15A/GADD34 in different cell population and infection conditions; and (D) violin plot representing the expression of PPP1R15A/GADD34 the whole PBMC population under uninfected, bystander and infected conditions.

## MATERIALS AND METHODS

### Cells and viruses

VERO-E6 (ATCC® CRL-1586™) A549 (ATCC® CCL-185™) and derived cell lines were grown at 37° C/5% CO2 in DMEM-F12 medium supplemented with 1 U/ml penicillin /streptomycin and 5% foetal bovine serum. C6/36 cells (ATCC® CRL-1660 were grown at 28° C in Leibovitz’s L-15 medium supplemented with 1 U/ml penicillin /streptomycin and 5% fetal bovine serum. IFNAR1 knockout cell line has been previously described (70, 71). PKR^-/-^ and DKO (IFNAR^-/-^/PKR^-/-^) were produced with a pair of sgRNA guides (PKR1-FWD: CACCGATTATGAACAGTGTGCATCG; PKR1-REV: AAACCGATGCACACTGTTCATAATC; PKR2-FWD:CACCGAAACAGTTCTTCGTTGCTTA; PKR2-REV: AAACTAAGCAACGAAGAACTGTTTC) cloned into pSpCas9(BB)-2A-Puro (PX459) vector (Addgene) according to developer’s protocol (72). Clonal populations of new cell lines were isolated and tested to confirm gene deletion (Supplementary figure 2). Experiments were carried out with the 1B6 clone of PKR^-/-^ cells and the 2B6 clone of DKO cells.

DENV-4 virus, TVP-360 strain (73) and ZIKV, strain BR 2015/15261 (74) were kindly provided by Dr Claudia N. Duarte dos Santos (Fiocruz-PR, Brazil). Viral stocks were produced in C6/36 cells and all viral titrations were performed in VERO cells by plaque-forming units (PFU) or focus-forming units (FFU).

### Poly (I:C)Transfection

Cell stimulation with poly (I:C) (“polyinosinic: polycytidylic acid”) was performed with Lipofectamine 3000 (Invitrogen) following manufacturer’s instructions for A459 cells at the proportion of 1 ug of poly(I:C) in 1 ml of medium in a 12-well plate.

### Western-blots

Samples were lysed on ice with RIPA buffer (150 mM NaCl; 50 mM Tris-HCl pH8; 1% NP40; 0.1% SDS; 0.5% sodium deoxycholate) supplemented with cOmplete™ and PhosSTOP inhibitors™ (Roche) according to manufacturer’s recommendations. Protein extracts were heat-denatured in the loading buffer and resolved in SDS-PAGE. Antibodies used for immunoblot reactions were: PKR antibody (Santa Cruz Biotechnology, Inc, sc-707, dilution 1:100); phospho-PKR (Thr451) polyclonal antibody, Invitrogen, 44-668G, dilution 1:100); phospho-eIF2α (Ser51) antibody (Cell Signaling Technology, Inc., #9721, dilution 1:1000); GADD34 antibody (Proteintech, 10449-1-AP, dilution 1:500); α-tubulin (DM1A) mAb (Cell Signaling Technology, Inc., #3873, dilution 1:1000); STAT1 antibody (AbCam, ab3987, dilution 1:1000); phospho-STAT1(Y701) M135 (Abcam, ab29045, dilution 1:1000); IFIT1 polyclonal antibody (Thermo/Pierce, PA3-848, dilution 1:1000); ISG15 mouse mAb (R&D System, MAB4845, dilution 1:1000); anti-mouse IgG, HRP-linked Antibody (Cell Signaling Technology, Inc., #7076, dilution 1:5000); anti-rabbit IgG, HRP-linked Antibody, Cell Signaling Technology, Inc., #7074, dilution 1:5000). The imaging of blots was carried out on the Chemidoc MP (Bio-Rad).

### Flow cytometry

Cells were fixed in BD-Phosflow Lyse/Fix buffer. Primary and secondary antibodies were diluted in permeabilization buffer (0.1% saponin; 1% fetal bovine serum, in PBS) for staining. Between each step, samples were washed in blocking buffer (1% fetal bovine serum in PBS). Antibodies and dyes used for flow cytometry were: phospho-eIF2α (Ser51) antibody (Cell Signaling Technology, Inc., #3597, dilution 1:300); human monoclonal antibody DV 18.4 (75)(dilution 1:100); GADD34 antibody (Proteintech, 10449-1-AP, dilution 1:250 (IF/FACS); Zombie-NIR fixable viability dye (BioLegend, 423105, dilution 1:750); Goat anti-Rabbit IgG (H+L) Alexa Fluor 647 conjugated (Invitrogen, A-21245, dilution 1:500); Goat anti-Rabbit IgG (H+L) Alexa Fluor 488 conjugated (Invitrogen, A-11008, dilution 1:500); Goat anti-Human IgG (H+L) Alexa Fluor 488 conjugated (Invitrogen, A-11013, dilution 1:500). Data acquisition was performed with FACSVerse™ (BD) and data analysis with FlowJo software (BD).

### Immunofluorescence

Cells were grown and infected on 13 mm cover slips in 24-wells plates and incubated in liquid or semisolid medium according to experiment design. Fixation in 3% paraformaldehyde for 20 minutes and permeabilization with Triton X-100 0,5% in PBS for 5 minutes. Primary and secondary antibodies were diluted in blocking buffer (2% bovine serum albumin in PBS) for staining. Between each step, samples were washed in PBS. Antibodies and dyes used for immunofluorescence were: DAPI (Invitrogen); anti-dsRNA MAb J2 (SCICONS, 10010500, dilution 1:200); human monoclonal antibody DV 18.4 (75)(dilution 1:100); anti-puromycin 12D10 (Millipore, MABE343, dilution 1:20,000); phospho-eIF2α (Ser51) antibody (Cell Signaling Technology, Inc., #3597, dilution 1:300); Goat anti-Rabbit IgG (H+L) Alexa Fluor 647 conjugated (Invitrogen, A-21245, dilution 1:500); Goat anti-Rabbit IgG (H+L) Alexa Fluor 488 conjugated (Invitrogen, A-11008, dilution 1:500); Goat anti-Human IgG (H+L) Alexa Fluor 488 conjugated (Invitrogen, A-11013, dilution 1:500); Donkey Anti-Mouse IgG H&L Alexa Fluor 647 conjugated (Abcam, ab150107, dilution 1:500). Image acquisition was performed in the confocal microscope Leica DMI6000 B.

### RT-qPCR

RNA was extracted and purified with HiYield TotalRNA mini kit (RBC Real Biotech Corporation, YRB50) according to manufacturer’s instructions. Reverse transcription reaction was performed with MMLV "in house" kit from 1 μg of total RNA. Quantitative PCRs were performed with GoTaq ® qPCR Master mix (Promega) with the following primer sets: h18S (F: 5’-TAGAGGGACAAGTGGCGTTC-3’, R: 5’-CGCTGAGCCAGTCAGTGT-3’); hIFNL1 (F: 5’-TTCCAAGCCCACCACAACTG-3’, R: 5’-GAGTGACTCTTCCAAGGCGT-3’); hIFNB1 (F: 5’-AAACTCATGAGCAGTCTGCA-3’, R: 5’-AGGAGATCTTCAGTTTCGGAGG-3’); hTNFA (F: 5’-TCTTCTCGAACCCCGAGTGA-3’, R: 5’-CCTCTGATGGCACCACCAG-3’); hISG15 (F: 5’-TCCTGGTGAGGAATAACAAGGG-3’, R: 5’-TCAGCCAGAACAGGTCGTC-3’); hDDIT3 (F: 5’-GGAGCATCAGTCCCCCACTT-3’, R: 5’-TGTGGGATTGAGGGTCACATC-3’); hGADD34 (F:

5’-CTGGCTGGTGGAAGCAGTAA-3’, R: 5’-TATGGGGGATTGCCAGAGGA-3’); DENV4 vRNA (F: 5’-TTGTCCTAATGATGCTGGTCG-3’, R: 5’-TCCACCTGAGACTCCTTCCA-3’); ZIKV vRNA (F: 5’-CTGTGGCATGAACCCAATAG-3’, R: 5’-ATCCCATAGAGCACCACTCC-3’). Data acquisition with StepOne Plus real time PCR system (Applied Biosystems) and relative mRNA expression was calculated by 2^-ΔΔCT^ method.

### ZIKV Luciferase reporter vectors

The pSGDLuc vector previously described (76), was used as a template to design the *Renilla* luciferase and the ZIKV firefly luciferase reporters. The *Renilla* luciferase was amplified using the following oligonucleotides 5′-TCCGCCCAGTTCCGCCCATTCTCCGC-3′ and 5′-GCGCTCTAGATTATCTCGAGGTGTAGAAATAC-3′ and then cloned back into the pSGDLuc previously digested with *AvrII* and *XbaI* This will create a plasmid only expressing the *Renilla* luciferase gene under the T7 or SV40 promoter.

The first 194 nucleotides of the ZIKV genome were cloned first in the pSGD plasmid using the set of primers 5′-GCGCCTCGAGATAAGTTGTTGATCTGTGTGAATCAGAC-3′ and 5′-GCGCAGATCTGCCCCCAAAGGGGCTCACACGGGCTAC-3′ in order to introduce this fragment in frame with the 2A-firefly luciferase gene. Using this intermediate plasmid as a template, the ZIKV-2A-firefly luciferase gene was later amplified with oligonucleotides 5′-TCCGCCCAGTTCCGCCCATTCTCCGC-3′and 5′-GATTCACACAGATCAACAACTccctatagtgagtc to introduce the T7 promoter upstream of the ZIKV coding region; and 5′-GCTTTACTGGGGCTACGATCTTTTGC-3′ and 5′-gactcactatagggAGTTGTTGATCTGTGTGAATC-3′ to amplify the ZIKV region. This was cloned back into the pSGD plasmid previously digested with *AvrII* and *BglII* . This will create a plasmid expressing the ZIKV-2A-firefly luciferase fused gene under the T7 or SV40 promoter.

### Luciferase assays

Cells were transfected using Lipofectamine 3000 (Invitrogen) with firefly and *Renilla Luciferase* plasmids at the proportion of 98 ng and 2 ng respectively for every 2x10^4^ cells in a 96-well plate. For cell lysis and luminescence reaction, Dual-Luciferase® Reporter Assay System (Promega) was used following the manufacturer’s instructions.

### Single-cell RNA sequencing analysis

Processed, publicly available single-cell RNA-seq data are available through the GEO accession number GSE110496 and GSE116672. We downloaded processed single-cell data and metadata from the supplementary information from the respective publication (55, 61).

Then, we used CellRouter (77) to perform quality control, normalisation, analysis and visualise the expression of selected genes with log2 fold changes > 0.15.

## AUTHOR CONTRIBUTIONS

Conceptualization, T.R.J. and D.S.M.; Methodology, T.R.J. and D.S.M.; Formal Analysis and Investigation, T.R.J.; *In silico analysis*, E.L.R., E.G.K., G.F.R.L.; Resources, D.S.M., B.F., N.I., T.S; Writing-Original Draft, T.R.J. and D.S.M., Writing-Review and editing, T.R.J., B.F., N.I., T.S and D.S.M., Visualization, T.R.J. and D.S.M., Supervision, D.S.M., Project Administration and Funding Acquisition, D.S.M.

## DECLARATION OF INTERESTS

The authors declare no competing interests.

## ACKNOWLEDGEMENTS

This work was supported by Comissão de Aperfeiçoamento de Pessoal de Nível Superior (CAPES) Computational Biology programme (23038.010048/2013-27), Conselho Nacional de Desenvolvimento Científico e Tecnológico (CNPq) (1 311406/2017-3 and 407609/2018-0) and the Academy of Medical Sciences/U.K. (NAF004/1005) to D.S.M., Conselho Nacional de Desenvolvimento Científico e Tecnológico (CNPq) (41907/2017-7) to T.R.J., BBSRC grant BB/S001336/1 and Wellcome Trust grant 201946/Z/16/Z to B.J.F; BBSRC grants BB/S007350/2 and BB/XO11046/1 and Wellcome Trust and Royal Society Sir Henry Dale Fellowship (202471/Z/16/A) to T.R.S; and an Isaac Newton Trust Grant (18.40r), a Royal Society Research Grant (RGS\R1\191137) and an Isaac Newton Trust/Wellcome Trust ISSF/University of Cambridge Joint Research Grant to NI.

Multi-User Laboratory in Biological Studies (LAMEB) and LCME at UFSC.

## Notes

### Competing Interest Statement

The authors have declared no competing interest.

### Summary of Updates

This version has updates in writing and new data.

